# Direct conversion of human fibroblasts into osteoblasts and osteocytes with small molecules and a single factor, Runx2

**DOI:** 10.1101/127480

**Authors:** Yanjiao Li, YaoLong Wang, Juehua Yu, Zhaoxia Ma, Qiong Bai, Xingfei Wu, Pengfei Bao, Lirong Li, Daiping Ma, Jinxue Liu, Change Liu, Fangyun Chen, Min Hu

## Abstract

Human osteoblasts can be induced from somatic cells by introducing defined factors, however, the strategy limits cells therapeutic applications for its multi-factor and complicated genetic manipulations that may bring uncertainty into the genome. Another important cell type in bone metabolism, osteocytes, which play a central role in regulating the dynamic nature of bone in all its diverse functions, have not been obtained from transdifferetiation so far. Herein, we have established procedures to convert human fibroblast directly into osteocyte-like and osteoblast-like cells using a single transcription factor, Runx2 and chemical cocktails by activating Wnt and cAMP/PKA pathways. These induced osteoblast-like cells express osteogenic markers and generate mineralized nodule deposition. A good performance of bone formation from these cells was observed in subcutaneous site of mouse at 4 weeks post-transplantation. Moreover, further studies convert human fibroblasts into osteocyte-like cells by orchestrating timing of the aforementioned chemical cocktails exposure. These osteocyte-like cells express osteocyte-specific markers and display characteristic morphology features of osteocytes. In summary, this study provides a promising strategy for cell-based therapy in bone regenerative medicine by direct reprogramming of fibroblasts into osteocytes and osteoblasts.

## Introduction

The advance of induced pluripotent cells (iPSCs) from reprogramming somatic cells and transdifferentiation from one adult somatic tissue to another type have provided a promising cell source for application in regenerative medicine therapy (Li et al., 2005; Takahashi et al., 2006; Takahashi et al., 2007; Kriks et al., 2011). The breaking discovery of iPSCs upon overexpression of four transcription factors (OCT4, KLF4, SOX2 and c-MYC) or by combination of small molecules has shed light on interrogating the mechanisms of cell fate reshaping (Takahashi et al., 2006; Takahashi et al., 2007; Bar-Nur et al., 2014). Other cell fates have been stimulated to apply sets of lineage-instructive transcriptional factors (TFs) that may convert cells from a distinct lineage to another somatic cell type, also known as direct reprogramming or transdifferentiation, without passing through an intermediate pluripotent stage. Advances in effective cell-fate switching from fibroblasts into neurons, cardiomyocytes, ECs, and hepatocytes have been achieved (Ieda et al., 2010; Ginsberg et al., 2012; Margariti et al., 2012; Huang et al., 2014; Zhou et al., 2014). However, these approaches for direct transdifferentiation between two lineage-specific cells employed multiple genetic factors. Such genetic manipulations may cast doubt on the utility of reprogrammed cells for their safety (Qian et al., 2012; Song et al., 2012). Therefore, it is urgently needed to explore transdifferentiation with minimal genetic manipulation in a chemically defined setting. Especially, new strategies to induce fibroblasts into different cell lineages by replacing the destination cells reprogramming TFs with small molecules could achieve comparable transdifferentiation efficiency (Hou et al., 2013; Pennarossa et al., 2013; Sayed et al., 2015).

Osteocyte is the most abundant cell type in mature bone and play a crucial role in regulating the dynamic nature of bone (Schaffler et al., 2012), which serves as a sensory network to respond to the effects of external influences. Osteocytes are derived from a subpopulation of committed osteoblasts which become buried in bone matrix to transform to multiple dendritic cells (Dallas et al., 2010). During bone formation, the osteoblasts can either become embedded in bone as osteocytes, or become inactive osteoblasts (bone lining cells) (Manolagas et al., 2000), whose fate is dependent on the bone signaling (Franz-Odendaal et al., 2006). Meanwhile, osteoblasts serve as a central role in bone formation to produce a calcium and phosphate-based mineral that is deposited into the mineralized matrix of bone (Neve et al., 2011). A functional osteoblast decline may cause osteolytic pathological conditions such as osteoporosis (Rachner et al., 2011). Given the central role of osteocytes and osteoblasts in bone and system physiology, tremendous efforts have been spent to obtain these two kinds of cells *in vitro* and to understand the transition mechanism for cell-based therapies on patients with osteoporosis (Yamamoto et al., 2015; Mattinzoli et al., 2012; Krishnan et al., 2010).

Recent achievements in osteoblasts induction from human fibroblasts have been reported by genetically introducing osteoblast-specific transcription factors (Runx2, Osterix) and reprogramming factors (Oct4, L-Myc) (Yamamoto et al., 2015). However, the multiple and complicated ectopic transgenetic expressions inevitably harbor safety, efficiency and other technical issues. On the other hand, there have been no reports on the induction of human osteocytes using a fibroblast source from direct reprogramming. Thus, a substitution of ectopic TFs by treating with small molecules should be a desirable alternative to convert directly somatic cells to ostoblast or osteocyte cells.

Here, we report a new strategy that combines the treatment of dexamethasone and a defined small-molecule cocktail consisting of CHIR99021 (GSK3 inhibitor) and forskolin (adenylyl cyclase activator) for osteoblast-osteocyte transdifferentiation. These compounds enabled the direct conversion of human fibroblasts into osteoblasts with only one TF, Runx2. We also tested the capability of the converted osteoblast-like cells (iOBs) for bone forming *in vivo*. Moreover, these chemical cocktails could induce osteocytes from fibroblasts without any additional differentiation conditions. In summary, our study has established a means of obtaining a source of autologous osteoblasts and osteocytes for bone tissue engineering.

## Results

### Induction of osteoblasts from human fibroblasts by the combination of a defined factor and small molecules

Several lines of evidence have supported the important role of osteogenic transcription factor Runx2 during bone development, which could effectively promote osteoblastic differentiation (Ducy et al., 1997; Karsenty et al., 1999; Komori et al., 1997). Transduction with Runx2 induced osteogenic gene expression and mineralized matrix deposition in rat fibroblasts (Phillips et al., 2007). To determine whether Runx2 is critical for fibroblast-osteoblastic cell conversion, we introduced Runx2 expression in primary human fibroblast cells from an 8-year boy. More than 40% Runx2-transduced cells were obtained after 3 days of infection (Supplementary information, Figure S1). Moreover, many proof-of-principle reports have demonstrated a crucial role for Wnt/beta-catenin signaling in the early stage of bone formation by stimulation of Runx2 gene expression (Canalis et al., 2005; Gaur et al., 2005; Westendorf et al., 2004). Wnt pathways have been shown to direct transdifferentiation of murine non-osteogenic cells into osteoblasts by inducing epigenetic modifications (Cho et al., 2014). Furthermore, given that cAMP/PKA signaling also plays a prominent role in osteogenesis (Kang et al., 2004; Zhao et al., 2007), we performed treatment of Runx2 transducted fibroblast cells with different combinations of small molecules CHIR99021 (CH) and forskolin (F) (involved in Wnt and PKA pathways respectively), to test their ability to transdifferentiate into osteoblast-like cells under dexamethasone treatment (Figure 1A). The Runx2 transducted cells (3 days after transduction, Figure 1A) were cultured in four conditions: DEX (dexamethasone), DEX/F (dexamethasone and forskolin), DEX/CH (dexamethasone and CHIR99021) and DEX/F/CH (dexamethasone, forskolin and CHIR99021) until day 25, and then analyzed the temporal gene expression profiling of osteoblastic markers including mRNA expression of alkaline phosphatase (ALP), osteocalcin (OCN), collagen type I a 1 (COL1A1), osteonectin (ON), osteopontin (OPN) and bone sialoprotein II (BSP-II) on day 25 (Figure 1B). Transdifferentiated osteoblast-like cell identity was characterized by comparing the gene expression patterns of known osteoblastic genes to the fibroblast cells undergoing with basic medium plus dexamethasone alone (DEX). The results showed a significant up-regulation of known osteoblastic genes, especially COL1A1 and ON through the combination treatment of CHIR and forskolin (DEX/F/CH). Noticeably, both COL1A1 and ON increased moderately in DEX/CH but remained the same in DEX/F, indicating that CHIR itself could induce osteoblastic genes and forskolin enhanced the expression only in presence of CHIR. And similar trend was observed in ALP expression. DEX/F moderately induced upregulation of OCN mRNA to the same extent of DEX/F/CH whereas DEX/CH exhibited similar OCN level compared to control (DEX), indicating CHIR has no effect in instigating OCN expression (1B). For BSP and OPN, either CH or F has any effect on their expression given that all conditions show the same mRNA levels relative to control. ALP staining confirmed the same results as real-time RT-PCR (Figure 1C).

**Figure 1.**
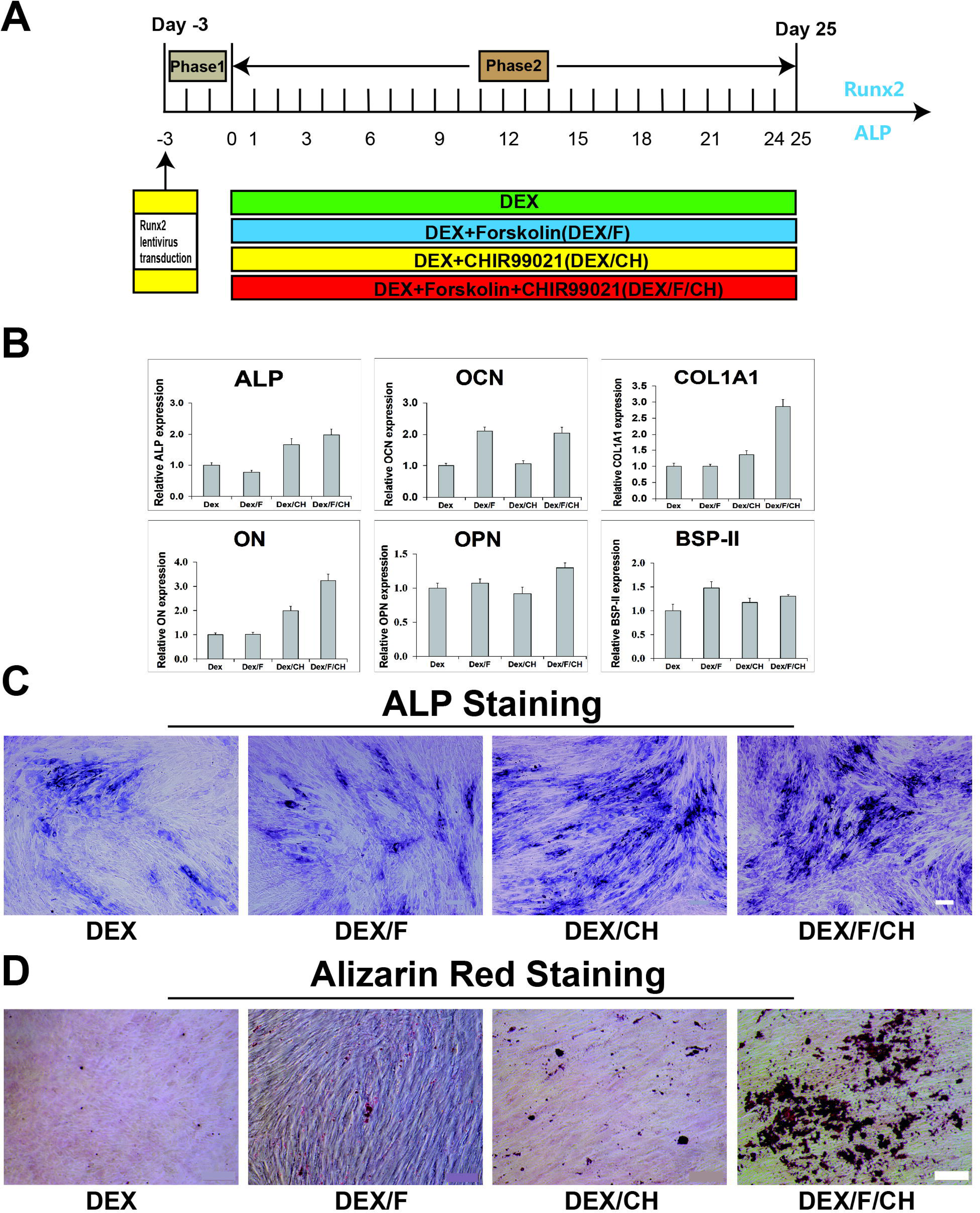
Induction of osteoblasts from human fibroblasts transduced by some combinations of small molecules and transduction with Runx2. **(A)** Diagram of culture conditions for induction of osteoblasts from human fibroblasts. Generally, initial fibroblasts were plated the day before lentivirus vectors infection and this day when cells were transduced with Runx2 is termed as “Day-3”. 3 days later, cells were transferred into DMEM with different combinations of small molecules in phase2. After culturing for 25 days, related markers for osteoblasts were detected. **(B)** Gene expression analysis for key osteoblastic cell markers on day25 by quantitative RT-PCR on day25. Significance compared to ‘DEX’. *P*<0.001 (*Dunnett’s test*). **(C)** ALP staining at day 25. **(D)** Alizarin Red staining at day 25 for mineral deposits(red). Scale bars, 100 μm for **C**, 200 μm for **D**.

We then examined the capability of calcium deposition in the four cultures by Alizarin Red staining (Paul et al., 1983). It showed that DEX/F/CH treatment induced apparent calcium deposition on day25 (Figure 1D). Therefore, the most remarkable changes of osteoblast-like cells were elicited by the combination of DEX/F/CH. And the DEX/F/CH induced osteoblast-like cells (iOBs) were tested in the subsequent experiment.

### Conversion efficiency of induced osteoblasts from human fibroblasts

To examine the efficiency of conversion from fibroblasts to iOBs, cells were dual-immunostained with Runx2 and Osterix on day 25 (Figure 2A). Osterix is another critical regulator of osteoblast differentiation, and is an indicator for osteoblast. Immunostaining of cells indicated that iOBs produced Osterix and Runx2 double-positive cells on day 25 (Figure 2A and 2B). The proportion of the Runx2-positive or Osterix-positive cells in total cells were 25.6±2.1% vs. 22.1±1.8% respectively under DEX/F/CH condition (Figure 2B). However, it was obvious that the enrichment of Osterix^+^ cells in Runx2-expressing cells, and the ratio of Osterix/Runx2 was up to 86.1±2.4 % in all Runx2^+^ cells (Figure 2B). The temporal decrease in Runx2 expression on day 25 may be a result of the non-transduced fibroblastic population proliferating faster than Runx2-expressing cells or cell necrosis by virus infection due to mass transport limitations.

**Figure 2.**
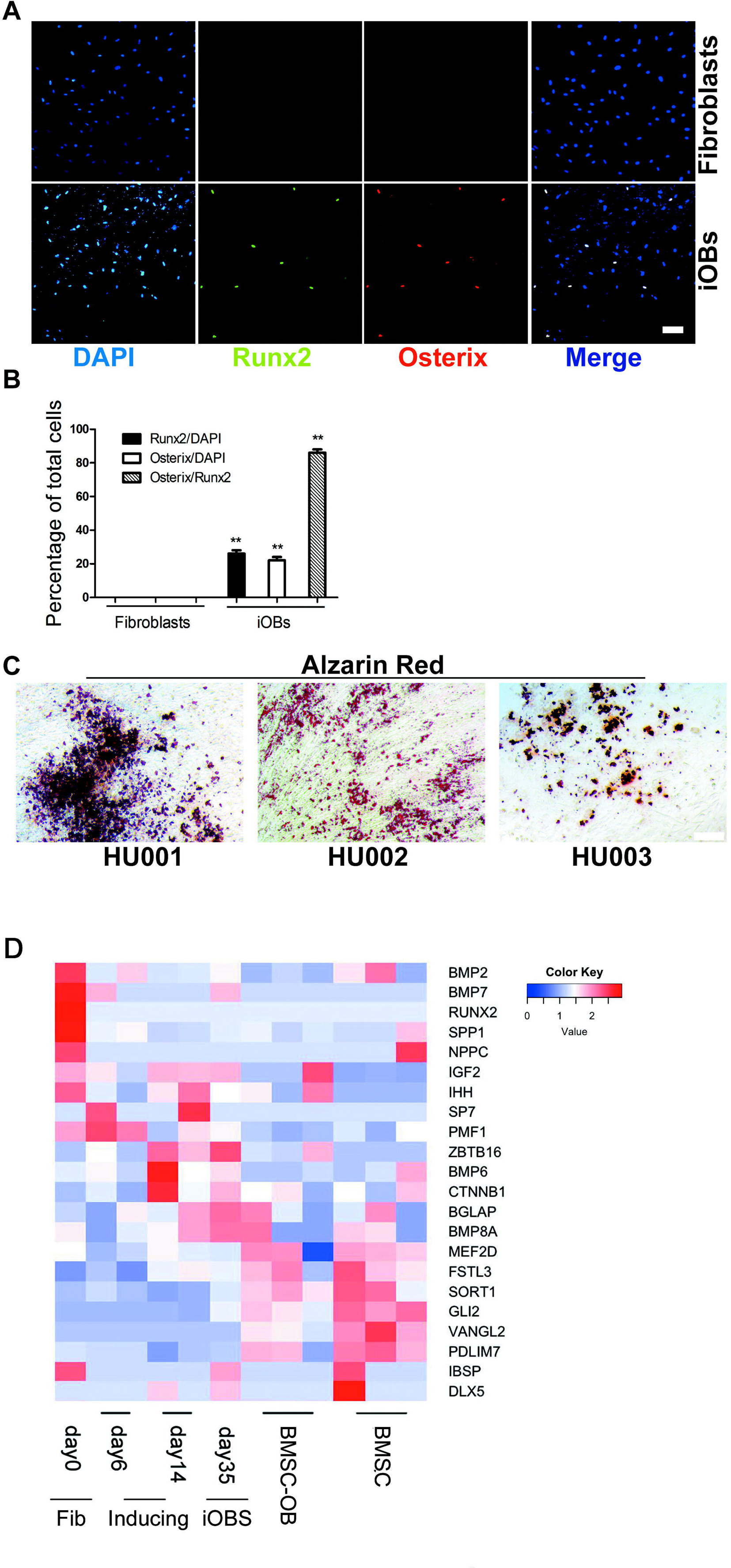
Conversion efficiency analysis of induced osteoblasts by DEX/F/CH treatment. **(A)** Immunocytochemistry at day 25 for co-expression of Runx2 (green) with Osterix (red) in fibroblasts (upper panels) and in iOBs (lower panels). Nuclei were counterstained with DAPI. The induced osteoblasts under DEX/F/CH conditions is termed as “iOBs”. **(B)** Quantification of the data presented in A to test the proportion of osterix producers. mean±s.e.m., n=3 (independent experiments), \*\**P*<0.01. **(C)** Calcium deposition in other three human fibroblasts derived osteoblasts by Alizarin Red staining. Scale bars represent 100 μm. **(D)** Heatmap showing osteogenic gene expression during reprogramming as compared to BMSCs and BMSC-derived osteoblasts.

To rule out the possibility that the induction of transdifferentiation could be due to loosen-up of the genomic structure by lentiviral insertion, we perfomed the four conditions (DEX, DEX/F, DEX/CH and DEX/F/CH) on fibroblasts transduced with empty vector with only GFP reporter. Runx2 and ALP were not induced after 25 days of culture in any culture indicating that Runx2 transduction is critical for following small molecule reprogramming into iOBs (Supplementary information, Figure S2).

To investigate whether iOBs could be derived from fibroblasts of another origin, and exclude off-target events, we transduced primary fibroblasts from three individual foreskin biopsies (HU001-HU003). All three primary human foreskin fibroblasts with Runx2 transduction by DEX/F/CH treatment evoked massive production of calcium deposition on day 25 (Figure 2C).

The heat map analysis showed that induced osteoblasts progressively expressed osteogenic gene. The expression pattern of iOBs was similar to that of BMSC-derived osteoblasts but distinguished from that of initial fibroblasts(Figure 2D).

To exclude the possibility that iOBs could be derived from contaminating MSCs (mesenchymal stem cells) mixed in fibroblast population, we cultured the fibroblasts in osteogenic and adipogenic differentiation media (Wang et al., 2009). No osteoblasts and adipocytes were produced from the fibroblasts, whereas human MSCs differentiated into these lineage-specific cells as a positive control (Supplementary information, Figure S3). The results clearly indicated that the iOBs were not arisen from MSCs contaminated in fibroblast population.

### iOBs are able to form bone tissue in vivo after subcutaneous transplantation in mice

We wanted to next investigate whether the osteogenic cells derived from Runx2-transduced fibroblasts could form viable bone tissue *in vivo*. Because of the induced osteogenesis, cells may quickly exited the cell cycle on day 25, we postulated the dissociation into single cells may cause cell death. Therefore, the cells converted in DEX/F/CH culture were digested into small clusters (by collagenase IV) on day 25 and mixed with matrigel before being transplanted into a subcutaneous site on the right back (3x10^5^ cells in 100 μl per site per animal) in six B6SJLF1 mice without prior cell purification. The control includes a matrigel engrafted alone without cells on the left. These mice were immuno-suppressed by ciclosporin everyday. The transplants were retrieved after 4 weeks post transplantation in all 6 mice. Histologically proven bone was formed in the cell graft site in four out of six donors analyzed while none formed with matrigel only controls in all six mice (Supplementary information, Figure S4A, B). No teratoma structures were observed in the cell implants or in the control site in all six mice. The size of implants was less than 1mm in diameter after 4 weeks (Figure 3A, B). Polarized light microscopy of the deposited matrix showed the lamellar structure of new bone, which indicated that bone tissue had been remodeled by osteoblasts and osteoclasts. Under ultraviolet light, the intense green fluorescence illustrated new bone formation consistent with highly mineralized matrix (Figure 3C). Histological examination of the explanted grafts by Alizarin Red or Basic fuchsin staining also showed that mineralized bone matrix generation, new bone formation. And mineralizing osteoblasts could be detected lining on the side of the mineralized matrix (Figure 3D, E).

**Figure 3.**
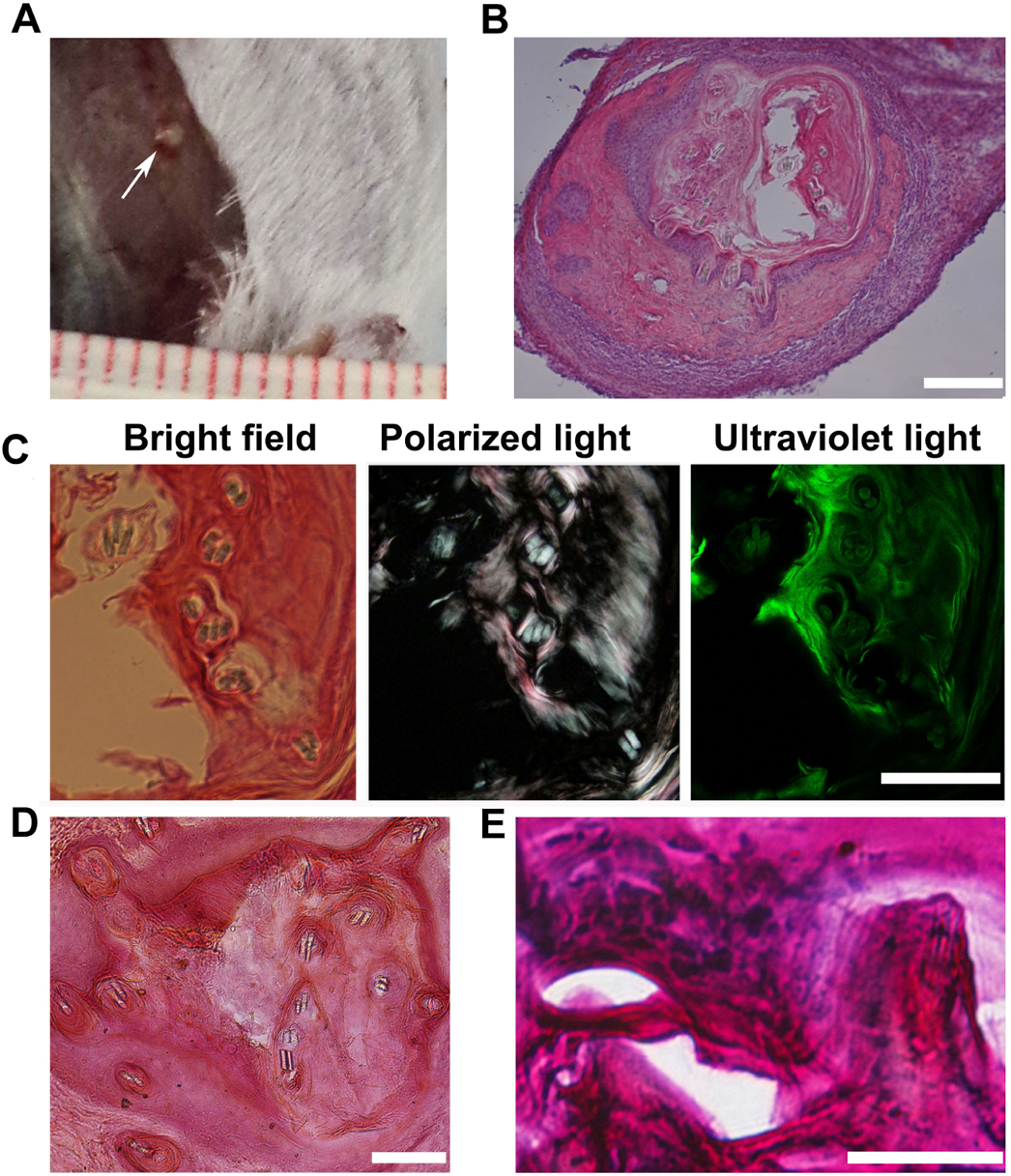
*In vivo* the bone-forming capacity of induced osteoblastic cells in mouse hosts. **(A)** The grafts (white arrow) were harvested as late as 4weeks post-implantation, which were grown on the surface of muscle under the skin. **(B)** A representative histological section stained with H&E showing the implanted cells to generate new tissue mixed with matrigel. **(C)** under bright field, polarized, or ultraviolet light the H&E-stained slides observed bone formation. Under polarized light, collagen bundles in bone matrix organized in parallel patterns observed implying the lamellar structure of new bone formation. Under ultraviolet light, intense green fluorescence demonstrated highly mineralized bone matrix. **(D)** Histological sections stained with Alizarin Red showing the mineralized matrix(red). **(E)** Basic fuchsin and methylene blue stained slides visualized the new bone formation(red) and lining osteoblasts along the bone matrix. Scale bars,100 μm for **B-D**, 50 μm for **E**.

The method was also applied to generate iOBs from rhesus monkey and tree shew fibroblasts (data not shown).

### Sustained expression of critical regulators for osteogenic differentiation by the chemical cocktails

To further characterize the process of osteogenic conversion by different chemical combinations, we analyzed the critical regulators in osteoblast differentiation. Runx2 is a master regulator for osteogenic development by enhancing the transcription of all major osteoblast-related genes. The expression of transduced exogenetic Runx2 was on the obvious decline on day10 (Figure 4A) with a maximum effect from day3 to day14 monitored by GFP green fluorescence in the four groups through the quantification of GFP positive cells in total cells (Figure 4B). Treatment with DEX or DEX/F showed low expression of Runx2 by day 14, but DEX/CH and DEX/F/CH were efficient at generating Runx2^+^ cells by gene expression analysis (Figure 4C) and immunocytochemistry (Figure 4D). This result implied that exogenous transduced Runx2 was silenced gradually (Figure 4B) and the endogenous Runx2 was activated (Figure 4C) in the processing of osteogenic induction before day14. Transient expression of Runx2 transgene was sufficient to induce cell-fate conversion from day3 to day14. Meanwhile, our data showed that up-regulation of endogenous Runx2 required activation of Wnt signaling using CHIR, alone or in combination with forskolin from day14 and forskolin had marginal effects whereas DEX/F maintained transgene Runx2 expression at high level until day12 (Figure 4B, C).

**Figure 4.**
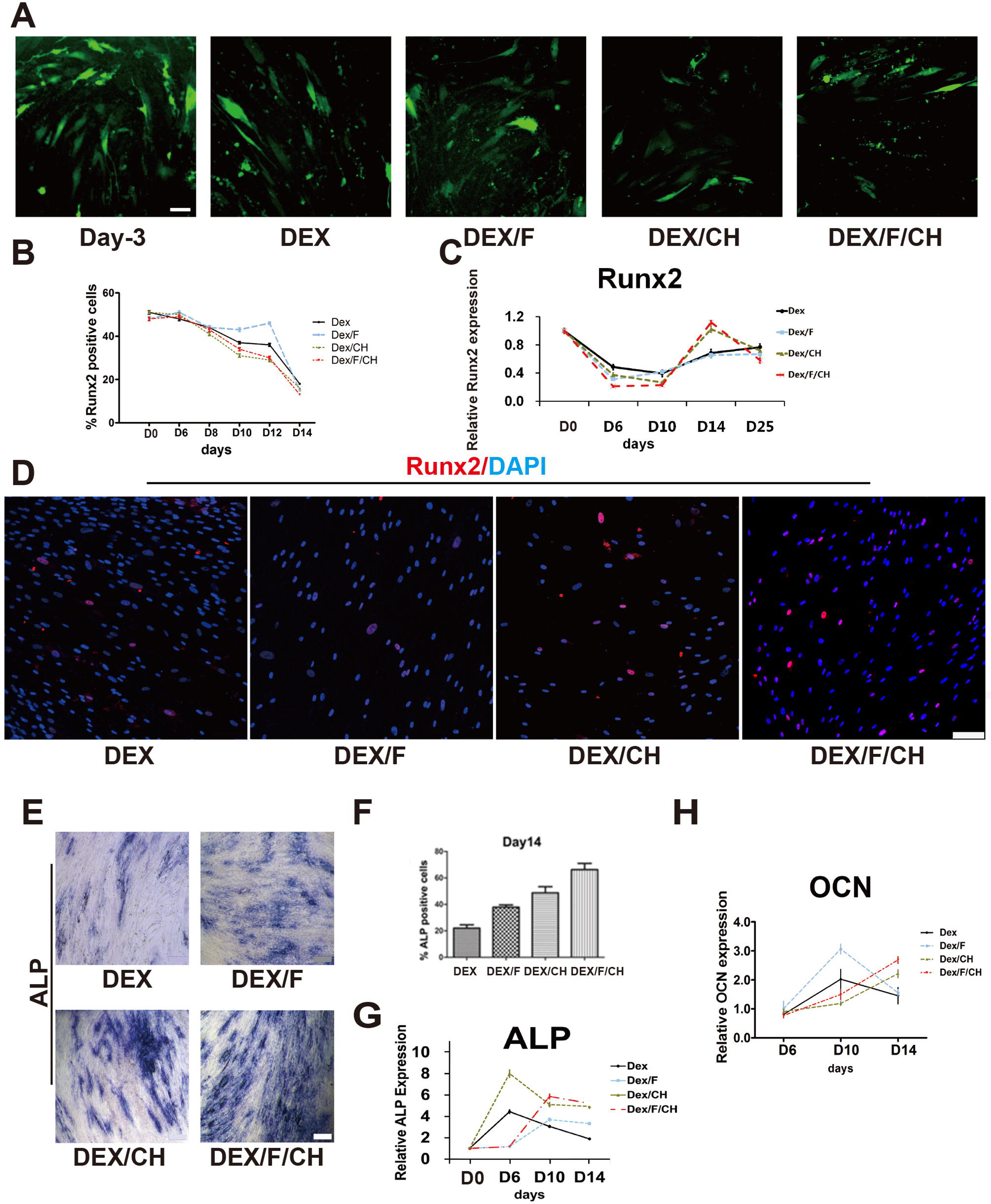
Characterization of the critical osteogenic regulators during cell-fate conversion by different chemical combinations. **(A)** The expression of transduced exogenetic Runx2 (green for GFP) in fibroblast infected by lentivirus vectors after 3 days on day0, and at day 10 under DEX, DEX/F, DEX/CH, DEX/F/CH respectively. Scale bars represent 75 μm. **(B)** Quantification of Runx2-GFP positive cells in total cells under four conditions above from day3 to day14. mean±s.e.m., n=3 (independent experiments): *P*<0.01. **(C)** Temporal gene expression analysis for *Runx2* from day0 to day25. Significance compared to ‘DEX’ at day0 (*Dunnett’s test*), P<0.001. **(D)** Representative images of induced osteoblast-like cells at day 14 stained for Runx2 antibody(red). Nuclei were stained with DAPI (blue). Scale bars represent 100 μm. **(E)** ALP staining at day 14 under above four conditions. Scale bars represent 200 μm. **(F)** Quantification of the data presented in **E**, analyzed to test ALP^+^ cells in total cells at day14. mean±s.e.m., n=3 (independent experiments): *P*<0.01. **(G)** Temporal gene expression analysis for *ALP* from day3 to day14. Significance compared to ‘DEX’ (*Dunnett’s test*), mean±s.e.m., n=3 (independent experiments) *P*<0.001. **(H)** Temporal gene expression analysis for *OCN* from day3 to day14. Significance compared to ‘DEX’ (*Dunnett’s test*), mean±s.e.m., n=3 (independent experiments) *P*<0.01.

Another key feature of skeleton developing is the enhanced expression of alkaline phosphatase (ALP), which is an early osteogenic marker. In this study, for the first 14 days, the efficiency of ALP induction was dependent on the treatment with the factor CHIR, forskolin, or the combination (Figure 4E). ALP was robustly induced in DEX/F/CH treated culture especially according to the quantification results on day14 (Figure 4F). Dexamethasone alone treatment could increase the ALP expression at day6, but was reduced since then (Figure 4G). Gene expression analysis revealed peak expression of ALP under DEX/CH condition, but ALP expressing was reduced gradually from day 6 to day 10 (Figure 4G), indicating CH is able to stimulate ALP expression at an early time point and keeps ALP at a reduced level later on. Forskolin first repressed ALP expression in DEX/F and DEX/F/CH, but released its repression from day 6 and enhanced ALP expression on top of CH in DEX/F/CH (Figure 4G). ALP mRNA levels were up-regulated by 3 orders of magnitude at day 10 compared to the cells at day 6 in DEX/F cultures, but were approximately 5 times at day 10 higher than at day 6 in DEX/F/CH consequently (Figure 4G).

OCN is one of characteristics for osteoblasts. Induction of OCN high expression required activation of PKA signaling using forskolin alone by day 10 in DEX/F culture, but the expression began to decline since then (Figure 4H). Treatment with CHIR or the combination of CHIR and forskolin showed repressed OCN first and increased OCN expressing from day 10 (Figure 4H). Therefore, these findings suggested that the timing of defined chemical cocktails exposure would take significant effects on osteogenic gene expression levels. With the presence of CH and F together in the medium, cells were kept in a primed state until day 6 and ready for boosting ALP and OCN from day 10-day14. Thus both molecules are required for accurate timing for osteoblastic transdifferentiation.

### Induction of osteocytes from human fibroblasts depends on the timing of small molecule exposure

Considering it is vital for lineage-specific cell reprogramming that the expression of master regulators are sustained and up-regulated at the beginning, we set out to examine whether orchestrating the timing of chemical cocktails exposure could take effects on osteogenic conversion from fibroblasts. Based on the aforementioned results, we performed a new two-stage strategy from day0 after Runx2 transduced instead of the one stage of DEX/F/CH (Figure 5A). The two stage was carried out by first addition of DEX/F from day0 to day11 and then adding DEX/F/CH after day11 termed as DEX/F-DEX/F/CH. Immunocytochemistry for GFP antibody staining showed that obvious morphological changes appeared on the induced cells which were maintained in DEX/F/CH for iOBs and DEX/F-DEX/F/CH compared to fibroblasts cultured in basic DMEM with 10%FBS on day25 (Figure 5B). The DEX/F/CH treated cells with transgene (termed iOBs in figure) displayed classical cubiodal appearance, typical of osteoblast. Addition of DEX/F-DEX/F/CH (termed as iOCs in figure) changed the morphology of the Runx2 transfected cells to have ramifications extended from the cell body which was the feature of osteocytes (Figure 5B). The remarkable phenotypic change was elicited by changing the timing of CHIR exposure from day11. Hereafter, we examined the characteristics of the induced osteocyte-like cells (iOCs) in the subsequent experiment.

**Figure 5.**
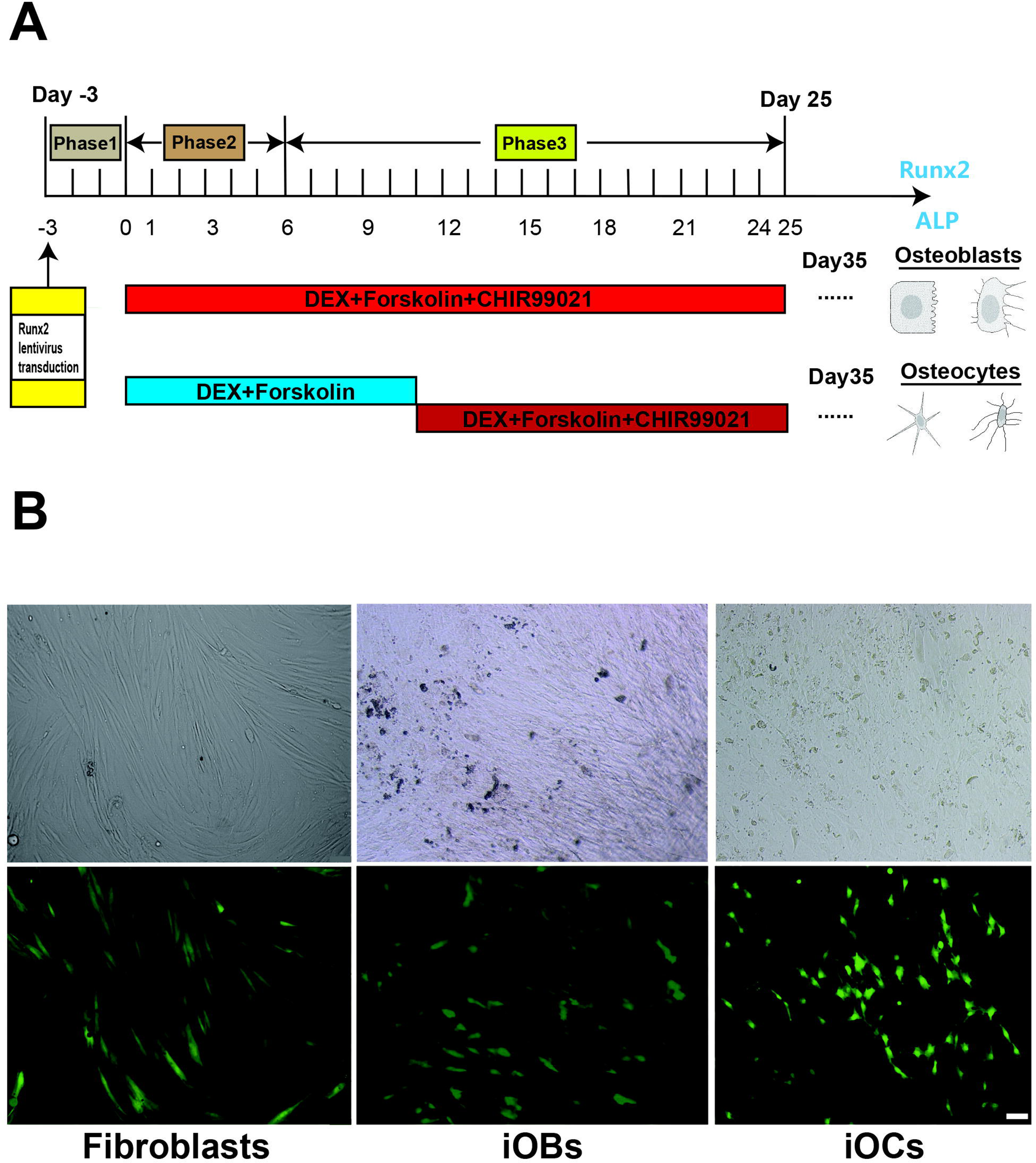
Induction of osteocyte-like cells from human fibroblasts transduced by the timing of defined small molecules cocktails exposure. **(A)** Diagram of culture conditions for induction of osteocytes from human fibroblasts. Initial fibroblasts were directed into osteocyte-like morphology including 3 phases. Induced osteocytes is termed as “iOCs”. **(B)** Representative images of converted cells stained for GFP on day25 under light microscopy (upper panels) and fluorescence (lower panels). Generally, Lentivirus vectors with Runx2-GFP insert were infected into human fibroblast at day-3. On day0, the cells with transgene were maintained in basic culture of DMEM including10%FBS, DEX/F/CH and DEX/F-DEX/F/CH respectively to direct fibroblasts, iOBs and iOCs correspondingly. Scal bar represents 75 μm.

### Cell morphology and differentially expressed genes of iOCs

We then sought to characterize the *in vitro* properties of iOCs under DEX/F-DEX/F/CH condition by day35. The fibroblasts cultured in basic medium of 10% FBS-DMEM showed elongated and spindle appearance, and the iOBs switched from fibroblast-like morphology to the retracted and stellate shape in DEX/F/CH (Figure 6A). But DEX/F-DEX/F/CH treated cells displayed a more elongated shape and dendrites which were more evident for interspersed ramified cellular elements at higher magnification (arrows) in iOCs (Figure 6A). ALP staining showed that its expression was declined dramatically in iOCs treated with DEX/F-DEX/F/CH as compared with iOBs incubated with DEX/F/CH (Figure 6B). Correspondingly, matrix mineralisation, as assessed by alizarin red staining, was most abundant in iOBs, whereas it was barely detectable in iOCs (Figure 6B).

**Figure 6.**
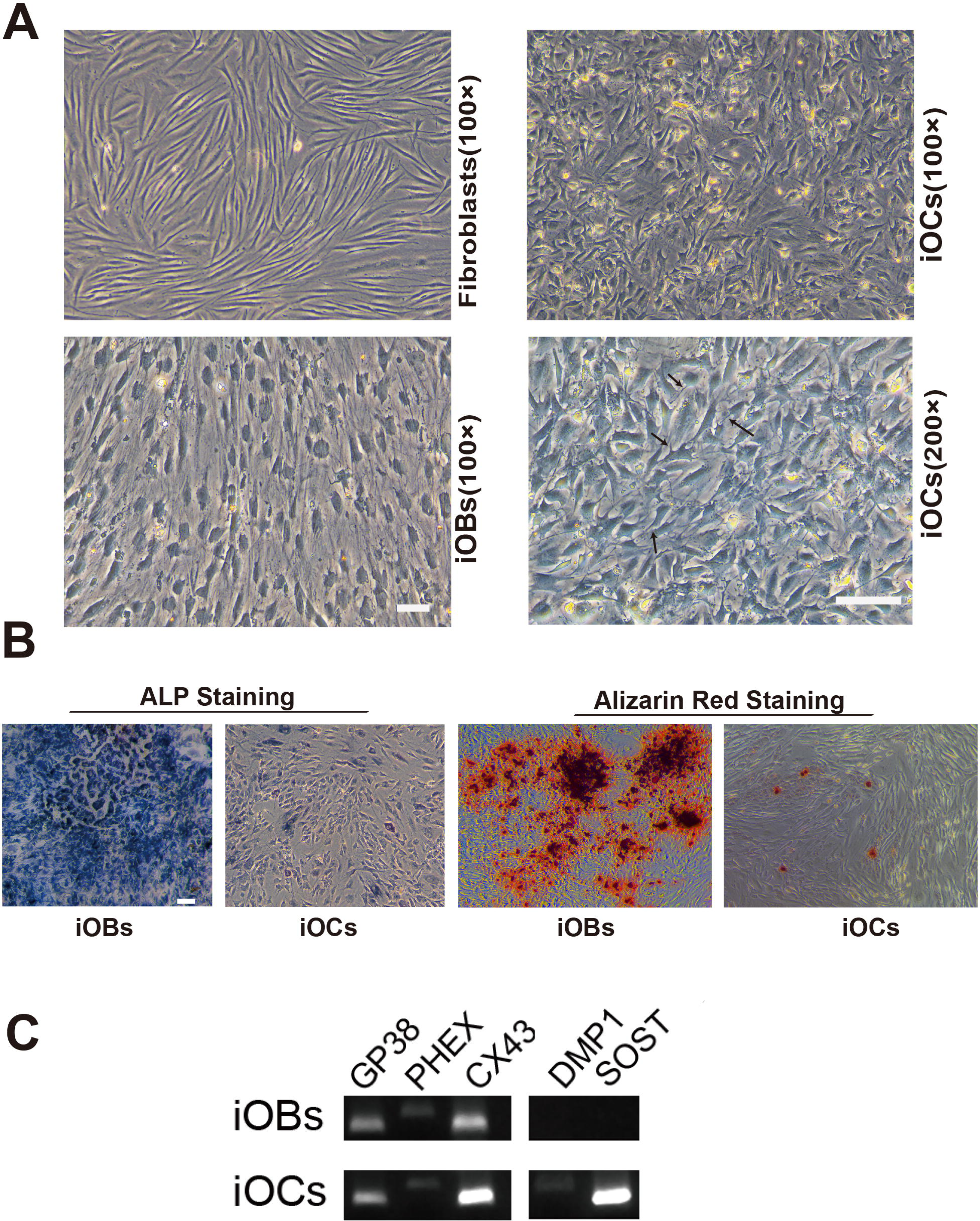
Characterization of morphology and gene expression of iOCs compared with iOBs. **(A)** Representative morphologies of fibroblasts, iOBs and iOCs on day35 under light microscopy. iOCs exhibit typical extended ramifications (arrows) compared with polygonal and cuboidal appearance of iOBs or elongated and spindle shape in fibroblasts. Scale bars represents 100 μm in 100X magnification, and in 200x magnification. **(B)** ALP staining for iOBs and iOCs conversion on day35. **(C)** Alizarin red staining for iOBs and iOCs conversion on day35. Scale bars represent 100 μm. **(D)** Gene expression for key osteocytes cell markers on day35 in iOBs and iOCs.

A panel of characteristic osteocyte markers in iOBs and iOCs were detected on day 35 such as podoplanin (also known as E11/GP38), phosphate-regulating gene with homology to endopeptidases on X chromosome (PHEX), connexin 43 (CX43), dentin matrix protein 1 (DMP1) and sclerostin (SOST) (Figure 6C). GP38, PHEX and CX43 were expressed in both iOBs and iOCs while DMP1and SOST were detected only in iOCs (Figure 6C). Our result of CX43 expression was in line with the reported evidences that CX43 played important role in function and survival of both osteoblasts and osteocytes. Given GP38 and PHEX expressed in iOBs population while no DMP1 and SOST observed in them, an “incomplete conversion” for osteocytes may exist under DEX/F/CH treatment. Accordingly, the majority of cells had cobblestone like features and the ramified cells were hardly observed in iOBs culture. But under DEX/F-DEX/F/CH procedure, the genes that were osteocyte specific or highly expressed in osteocytes have been all detected in iOCs population including DMP1 and SOST, implying that iOCs shared similar gene expression signature with osteocytes. Therefore, the iOC gene expression profiling needs to be further analyzed.

## Discussion

First of all, this is the first study demonstrating that osteocytic reprogramming can be achieved from human fibroblast cells. In this study, the iOCs are characterized by the formation of dendritic processes with a reduction in cytoplasmic volume and low level of mineralisation and alkaline phosphatase production (Dallas et al., 2010; Kato et al., 2001), while the iOBs are with the classical cuboidal appearance. Some authors have studied osteocytes *in vitro* by a 3D culture system where human osteoblast differentiation can be induced by matrix scaffolds (Boukhechba et al., 2009). Others found that or by specific compounds such as retinoic acid, mouse primary osteoblasts could be induced to osteocytes (Mattinzoli et al., 2012). The conversion process induced by our chemical cocktail strategy shows that these small molecules are enough to modulate osteocyte gene expression and promote induced transition from osteoblast, either by coordinating multiple signaling pathways or or by producing a subpopulation of osteoblasts destined for osteocytes. It is of note that the induced strategy presents a method for one-step osteocyte conversion without extra steps severing the process. However, the transitional stages and the underlying mechanism from fibroblasts to osteoblasts-osteocytes remain open for future research.

Second, we found that manipulation of the signal transduction milieu by combination of cAMP/PKA and Wnt pathways could enhance the level of ALP and endogenous expression of Runx2 to induce osteoblasts. As several lines of evidence have shown that these aforementioned signaling may interact and act at diverse stages of the osteogenesis program, i.e., Wnts stimulate Runx2 gene expression (Westendorf et al., 2004; Gaur et al., 2005; Canalis et al., 2004), and promote osteoblast precursor growth and some early events in osteoblast differentiation by changing ALP activity (Westendorf et al., 2004). Recent studies have shown that the activation of the cAMP/PKA pathway is crucial for bone development and can stimulate the osteogenic gene expression (Yang et al., 2004; Zhao et al., 2007). Therefore, the chemical cocktail combined with only one lineage-specific TF, Runx2 may be sufficient to up-regulate osteogenic gene expression specifically and erase fibroblast gene expression.

But it is interesting that once the cells were primed with DEX/F first followed by DEX/F/CH treatment, the transitional stages from fibroblasts to osteoblasts or osteocytes have been altered correspondingly. It is obvious that the timing of CHIR and forskolin exposure takes crucial effects on the osteoblasts or osteocytes conversion from fibroblasts. A proper fine-tuning of cAMP/PKA and Wnt pathways is critical for the two cell types induction from human fibroblasts by small molecules. Interestingly, we found that the timing of different chemical cocktails exposure in reprogramming may be as important as the combination of different small molecules itself. However, the different timing of chemicals exposure and the precise mechanisms underlying this strategy for induction of osteoblast–osteocyte cells remain to be further determined considering these signaling pathways and TFs working/interacting complexly.

Third, we also found that effective osteoblast reprogramming can be achieved from fibroblast cells using a single transcriptional factor enabled by small molecules. Compared with Dr. Yamamoto’s osteoblast direct conversion method by transducing four defined factors which are Runx2, Osterix, Oct4 and L-Myc (Yamamoto et al., 2015), our study is consistent with their finding that induced osteoblasts express endogenous Runx2, not requiring continuous expression of the exogenous genes to maintain their phenotype. Our findings show that the combination of CHIR and forskolin could replace the effects of genetic factor Oct4 and L-Myc to promote endogenous Runx2 expression, although the two TFs are important to cause endogenous Runx2 in Dr. Yamamoto’s report. On the other hand, although Dr. Yamamoto’s study demonstrates that Osterix plays a crucial role on osteoblasts “full conversion”, the endogenous Osterix expression could be achieved by the small molecules combination in our research without the TF transduced. This technology of small molecules reprogramming would provide an alternative for the study of osteogenic cell-fate switching. However, the precise mechanisms underlying the osteoblast reprogramming by combined exposure to Dex/F/CHIR and Runx2 transgene remain to be determined.

In conclusion, our study offers another strategy for the generation of individual specific osteoblasts and osteocytes which would provide desirable cell resources for disease modeling and for cell-based therapy against bone resorptive diseases.

## Materials and Methods

### Culture of human fibroblast cell

Primary fibroblasts were harvested from 8-year old male cells (passages 4-8) were expanded in growth medium consisting of DMEM (GIBCO), 10% fetal bovine serum (FBS) (Hyclone), 100 U/ml penicillin and 100 μg/ml streptomycin antibiotics (sigma) and all other cell culture supplements and reagents were acquired from Sigma. Other three fibroblasts (HU001, HU002, HU003) were obtained from three adult persons’ foreskin respectively.

### Lentiviral transduction

The Runx2 lentiviral vector was pCMV-VSV-G system to express the human Runx2 gene tagged with GFP, and the empty vector control lacked Runx2 insert. Plasmid DNA was purified from transformed Escherichia coli using plasmid DNA Purification kits (promega). Lentiviruses were packaged using Lipofectamine 2000 (Invitrogen) as described elsewhere and got a high titer virus. Human skin fibroblasts of passage 4-8 was transduced by CSC-Runx2-ires-GFP or empty CSC-ires-GFP lentivirus as control at 40–60% confluence maintained in 10% FBS DEME medium with 100U/ml penicillin and 100 μg/ml streptomycin antibiotics for three days at the beginning. Runx2-transduced cells were analyzed for transduction efficiency using quantification of GFP expression via flow cytometry (BD FCs Jazz cell sorter) after 72 hours.

### Osteoblastic and osteocytes cells induction

In osteogenic cells induction, we referred to ‘DEX’ treatment as exposure of cells to steroid hormone dexamethasone (10 nM, sigma) in the basic medium including 10% FBS (Hyclone), 100 U/ml penicillin (Sigma), 100 μg/ml streptomycin (Sigma), DMDM (Gibco).

In other groups, ‘DEX/F’ as exposure to dexamethasone (10nM) and forskolin (10 μM,Enzo), ‘DEX/CH’ as dexamethasone (10 nM) and CHIR99021(9 μM,Stegment) and ‘DEX/F/CH’ as the combination of dexamethasone (10 nM), forskolin (10 μM, Enzo) and CHIR99021 (9 μM, Selleck) in the basic medium from day0 to day25. And the cells at day25 were used for transplantation studies. For osteocytes induction, cells were maintained in DEX/F/CH till day35.

### Gene expression analyses

Total RNA was extracted during differentiation at day 3, 6, 10, 14 and 25 from each condition of DEX, DEX/F, DEX/CH and DEX/F/CH using an RNeasy kit (Qiagen). For qRT–PCR analyses, total RNA at each condition was reverse transcribed (Takara) and amplified material was detected using commercially SYBR green probes (Takara) with the data normalized to HPRT control. All results were from 3-5 technical replicates of 2–3 independent biological samples at each data point. Some of primers were the same as previously reported (Boukhechba et al., 2009; Phillips et al., 2007).

### Animal surgery

All animal procedures were carried out under the local Institutional Animal Care and Use committee (IACUC). Mice: A total number of 6 B6SJLF1 mice (20-35 g) were used in this study. They were anaesthetized with ketamine (90 mg/kg; Gutian, China). *In vivo* bone formation assays, a 1-2 cm skin incision was made to create a subcutaneous pocket. A total of 3x10^5^ cells mixed with matrigel (BD) in total 100 μl were placed into the pocket of right lumbar back in one mouse, and the contralateral side was injected with matrigel (BD) of same volume. After surgery, mice were put back to their original cages. The mice received injection in the subcutaneous site using ciclosporin (4 mg/kg) everyday. The mice were housed with free access to food and water.

### Tissue processing

Mice were injected with overdoses of pentobarbital intra-peritoneally (50 mg/kg) to result in the deep anaesthesia. The grafts were immediately post-fixed in 4% PFA then soaked in 20% sucrose solutions for 1-2 days. They were sectioned on a cryostat, after embedding in O.C.T. (Sakura-Finetek) at 18 μm serial sections coronally.

### Immunohistochemistry

Cells were fixed in 4% PFA and blocked with 2% BSA with 0.1% Triton. The graft sections were washed in cold PBS and processed similarly. ALP was stained by staining kit (Sidansai stem cell, China). Primary antibody of Runx2 (Santa cruz, sc-10758), GFP (Santa cruz, sc-5385) and Osterix (Santa cruz, sc-22538) diluted in 1% BSA and incubated according to the manufacturer recommendations. Alexa555 and Alexa 488 were conjugated secondary antibodies (Invitrogen) used with 4’ 9, 6-diamidino-2-phenylindole (DAPI) nuclear counterstain (Thermo Fisher) by a fluorescence microscope (Nikon). Graft tissue sections were stained with basic fuchsin, methylene blue, and H&E for new bone formation or stained with Alizarin Red for analyzing the mineralization matrix distribution by a light microscope (Nikon) (Siddappa et al., 2008; Lu et al., 2014).

### RNA-seq and heat map production

RNA-seq libraries were prepared from samples that passed QC according to the manufacturer’s protocol (Illumina). Sequencing to produce single-end 50-bp reads was then performed on an Illumina HiSeq 2500 instrument. Heat map generated from RNA-seq data reflecting gene expression values in different time points.

## Acknowledgments

We thank all members of the lab for sharing reagents and advice. This work was supported by the Natural Science Foundation of Kunming City, China (2014-04-A-S-01-3072).

## References

Bar-Nur, O., Brumbaugh, J., Verheul, C. et al. (2014). Small molecules facilitate rapid and synchronous iPSC generation. Nature methods 11, 1170–1176.

Boukhechba, F., Balaguer, T., Michiels, JF. et al. (2009). Human primary osteocyte differentiation in a 3D culture system. Journal of Bone and Mineral Research 24, 1927–1935.

Canalis, E., Deregowski, V., Pereira, RC. and Gazzerro, E. (2005). Signals that determine the fate of osteoblastic cells. J Endocrinol Invest 28, 3–7.

Cho, Y-D., Yoon, W-J., Kim, W-J. et al. (2014). Epigenetic modifications and canonical wingless/int-1 Class (WNT) signaling enable trans-differentiation of nonosteogenic cells into osteoblasts. Journal of Biological Chemistry 289, 20120–20128.

Dallas, SL. and Bonewald, LF. (2010). Dynamics of the transition from osteoblast to osteocyte. Annals of the New York Academy of Sciences 1192, 437–443.

Ducy, P., Zhang, R., Geoffroy, V., Ridall, AL. and Karsenty, G. (1997). Osf2/Cbfa1: a transcriptional activator of osteoblast differentiation. Cell 89, 747–754.

Franz-Odendaal, TA., Hall, BK. and Witten, PE. (2006). Buried alive: how osteoblasts become osteocytes. Developmental Dynamics 235, 176–190.

Gaur, T., Lengner, CJ., Hovhannisyan, H. et al. (2005). Canonical WNT signaling promotes osteogenesis by directly stimulating Runx2 gene expression. Journal of Biological Chemistry 280, 33132–33140.

Ginsberg, M., James, D., Ding, B-S. et al. (2012). Efficient direct reprogramming of mature amniotic cells into endothelial cells by ETS factors and TGFβ suppression. Cell 151, 559–575.

Hou, P., Li, Y., Zhang, X. et al. (2013). Pluripotent stem cells induced from mouse somatic cells by small-molecule compounds. Science 341, 651–654.

Huang, P., Zhang, L., Gao, Y. et al. (2014). Direct reprogramming of human fibroblasts to functional and expandable hepatocytes. Cell stem cell 14, 370–384.

Ieda, M., Fu, J-D., Delgado-Olguin, P. et al. (2010). Direct reprogramming of fibroblasts into functional cardiomyocytes by defined factors. Cell 142, 375–386.

Kang, Y. and Massague, J. (2004). Epithelial-mesenchymal transitions: twist in development and metastasis. Cell 118, 277–279.

Karsenty, G., Ducy, P., Starbuck, M. et al. (1999). Cbfa1 as a regulator of osteoblast differentiation and function. Bone 25, 107–108.

Kato, Y., Boskey, A., Spevak, L., Dallas, M., Hori, M. and Bonewald, L. (2001). Establishment of an Osteoid Preosteocyte-like Cell MLO-A5 That Spontaneously Mineralizes in Culture. Journal of Bone and Mineral Research 16, 1622–1633.

Komori, T., Yagi, H., Nomura, S. et al. (1997). Targeted disruption of Cbfa1 results in a complete lack of bone formation owing to maturational arrest of osteoblasts. Cell 89, 755–764.

Kriks, S., Shim, J-W., Piao, J. et al. (2011). Dopamine neurons derived from human ES cells efficiently engraft in animal models of Parkinson/’s disease. Nature 480, 547–551.

Krishnan, V., Dhurjati, R., Vogler, EA. and Mastro, AM. (2010). Osteogenesis in vitro: from pre-osteoblasts to osteocytes. In Vitro Cellular & Developmental Biology-Animal 46, 28–35.

Li, X-J., Du, Z-W., Zarnowska, ED. et al. (2005). Specification of motoneurons from human embryonic stem cells. Nature biotechnology 23, 215–221.

Lu, T., Hu, P., Su, X., Li, C., Ma, Y. and Guan, W. (2014). Isolation and characterization of mesenchymal stem cells derived from fetal bovine liver. Cell and tissue banking 15, 439–450.

Manolagas, SC. (2000). Birth and death of bone cells: basic regulatory mechanisms and implications for the pathogenesis and treatment of osteoporosis 1. Endocrine reviews 21, 115–1.

Margariti, A., Winkler, B., Karamariti, E. et al. (2012). Direct reprogramming of fibroblasts into endothelial cells capable of angiogenesis and reendothelialization in tissue-engineered vessels. Proceedings of the National Academy of Sciences 109, 13793–13798.

Mattinzoli, D., Messa, P., Corbelli, A. et al. (2012). A novel model of in vitro osteocytogenesis induced by retinoic acid treatment. Eur Cell Mater 24, 403–425.

Neve, A., Corrado, A. and Cantatore, FP. (2011). Osteoblast physiology in normal and pathological conditions. Cell and tissue research 343, 289–302.

Paic, F., Igwe, JC., Nori, R. et al. (2009). Identification of differentially expressed genes between osteoblasts and osteocytes. Bone 45, 682–692.

Paul, H., Reginato, AJ. and Schumacher, HR. (1983). Alizarin red S staining as a screening test to detect calcium compounds in synovial fluid. Arthritis & Rheumatism 26, 191.

Pennarossa, G., Maffei, S., Campagnol, M., Tarantini, L., Gandolfi, F. and Brevini, TA. (2013). Brief demethylation step allows the conversion of adult human skin fibroblasts into insulin-secreting cells. Proceedings of the National Academy of Sciences 110, 8948–8953.

Phillips, JE., Guldberg, RE. and García, AJ. (2007). Dermal fibroblasts genetically modified to express Runx2/Cbfa1 as a mineralizing cell source for bone tissue engineering. Tissue engineering 13, 2029–2040.

Qian, L., Huang, Y., Spencer, CI. et al. (2012). In vivo reprogramming of murine cardiac fibroblasts into induced cardiomyocytes. Nature 485, 593–598.

Rachner, TD., Khosla, S. and Hofbauer, LC. (2011). Osteoporosis: now and the future. The Lancet 377, 1276–1287.

Sayed, N., Wong, WT., Ospino, F. et al. (2015). Transdifferentiation of Human Fibroblasts to Endothelial Cells Role of Innate Immunity. Circulation 131, 300–309.

Schaffler, MB. and Kennedy, OD. (2012). Osteocyte signaling in bone. Current osteoporosis reports 10, 118–125.

Siddappa, R., Martens, A., Doorn, J. et al. (2008). cAMP/PKA pathway activation in human mesenchymal stem cells in vitro results in robust bone formation in vivo. Proceedings of the National Academy of Sciences 105, 7281–7286.

Song, K., Nam, Y-J., Luo, X. et al. (2012). Heart repair by reprogramming non-myocytes with cardiac transcription factors. Nature 485, 599–604.

Takahashi, K. and Yamanaka, S. (2006). Induction of pluripotent stem cells from mouse embryonic and adult fibroblast cultures by defined factors. Cell 126, 663–676.

Takahashi, Y., Roesch, MR., Stalnaker, TA. and Schoenbaum, G. (2007). Cocaine exposure shifts the balance of associative encoding from ventral to dorsolateral striatum. Frontiers in integrative neuroscience 1, 11.

Wang, X. and Moutsoglou, D. (2009). Osteogenic and Adipogenic Differentiation Potential of an Immortalized Fibroblast-like Cell Line Derived from Porcine Peripheral Blood. In Vitro Cellular & Developmental Biology – Animal 45, 584–591.

Westendorf, JJ., Kahler, RA. and Schroeder, TM. (2004). Wnt signaling in osteoblasts and bone diseases. Gene 341, 19–39.

Yamamoto, K., Kishida, T., Sato, Y. et al. (2015). Direct conversion of human fibroblasts into functional osteoblasts by defined factors. Proceedings of the National Academy of Sciences 112, 6152–6157.

Yang, X., Matsuda, K., Bialek, P. et al. (2004). ATF4 is a substrate of RSK2 and an essential regulator of osteoblast biology: implication for Coffin-Lowry syndrome. Cell 117, 387–398.

Zhao, Y. and Ding, S. (2007). A high-throughput siRNA library screen identifies osteogenic suppressors in human mesenchymal stem cells. Proceedings of the National Academy of Sciences 104, 9673–9678.

Zhou, D., Zhang, Z., He, L-M. et al. (2014). Conversion of fibroblasts to neural cells by p53 depletion. Cell reports 9, 2034–2042.

